# Unique Transcriptomic Cell Types of the Granular Retrosplenial Cortex are Preserved Across Mice and Rats Despite Dramatic Changes in Key Marker Genes

**DOI:** 10.1101/2024.09.17.613545

**Authors:** Isla A.W. Brooks, Izabela Jedrasiak-Cape, Chloe Rybicki-Kler, Tyler G. Ekins, Omar J. Ahmed

**Affiliations:** Dept. of Psychology, University of Michigan, Ann Arbor, MI 48109; Neuroscience Graduate Program, University of Michigan, Ann Arbor, MI 48109; Dept. of Biomedical Engineering, University of Michigan, Ann Arbor, MI 48109

**Author notes:** **LEAD CONTACT & CORRESPONDING AUTHOR:** Omar J. Ahmed *Mail:* Dept. of Psychology, 530 Church St. University of Michigan, Ann Arbor, MI 48109 *Email:.

## Abstract

The granular retrosplenial cortex (RSG) supports key functions ranging from memory consolidation to spatial navigation. The mouse RSG contains several cell types that are remarkably distinct from those found in other cortical regions. This includes the physiologically and transcriptomically unique low rheobase neuron that is the dominant cell-type in RSG layers 2/3 (L2/3 LR), as well as the similarly exclusive pyramidal cells that comprise much of RSG layer 5a (L5a RSG). While the functions of the RSG are extensively studied in both mice and rats, it remains unknown if the transcriptomically unique cell types of the mouse RSG are evolutionarily conserved in rats. Here, we show that mouse and rat RSG not only contain the same cell types, but key subtypes including the L2/3 LR and L5a RSG neurons are amplified in their representations in rats compared to mice. This preservation of cell types in male and female rats happens despite dramatic changes in key cell-type-specific marker genes, with the *Scnn1a* expression that selectively tags mouse L5a RSG neurons completely absent in rats. Important for Cre-driver line development, we identify alternative, cross-species genes that can be used to selectively target the cell types of the RSG in both mice and rats. Our results show that the unique cell types of the RSG are evolutionarily conserved across millions of years of evolution between mice and rats, but also emphasize stark species-specific differences in marker genes that need to be considered when making cell-type-specific transgenic lines of mice versus rats.

## INTRODUCTION

The retrosplenial cortex (RSC) is a densely interconnected node of the default mode network and plays an important role in memory, navigation and imagining oneself in the future (Alexander et al., 2023; Auger et al., 2012; Chang et al., 2020; Miller et al., 2019, 2021; Mitchell et al., 2018; Spiers & Maguire, 2006; Vann et al., 2009). Across both rodents and primates, damage to the RSC is associated with severe impairments of navigational abilities and memory (Alexander et al., 2023; Buckley & Mitchell, 2016; Hindley et al., 2014; J. H. Kim et al., 2007; Mitchell et al., 2018; A. J. D. Nelson et al., 2015; Wesierska et al., 2009). While mice and rats have both been used to study retrosplenial cells, circuits, and behavioral functions (Alexander, Robinson, et al., 2023; Balcerek et al., 2021; Mitchell et al., 2018; Todd et al., 2019; Vann et al., 2009; Wyss & Van Groen, 1992), few studies have directly compared retrosplenial properties across these two species (but see the comparison of mouse and rat retrosplenial brain rhythms in Ghosh et al., 2022). Mice *(Mus musculus)* and rats *(Rattus norvegicus)* evolutionarily diverged between 12-36 million years ago (Adkins et al., 2001; Nei et al., 2001; Springer et al., 2003), suggesting that a careful comparison of mouse and rat RSC would reveal key evolutionarily conserved features central to retrosplenial function.

Several *in vivo* studies of cellular spatial encoding patterns have been pioneered in rats, given their more stable behavior and comparatively easier implantation of multiple probes due to their larger skull size (Alexander et al., 2020; Alexander & Nitz, 2015, 2017; Chen et al., 1994; Jacob et al., 2017; Lozano et al., 2017; O’Keefe & Dostrovsky, 1971; Smith et al., 2018; Taube et al., 1990b, 1990a; Vedder et al., 2017). Examples of this extensive *in vivo* characterization of rat RSC neurons includes the discovery of egocentric boundary vector cells (Alexander et al., 2020), RSC axis cells (Jacob et al., 2017), cells sensitive to path turning direction (Alexander & Nitz, 2015), and cells encoding context-dependent spatial location (Miller et al., 2021). Fear conditioning experiments in rats have additionally identified a critical role of the RSC in trace fear extinction (Keene & Bucci, 2008b, 2008a; Kwapis et al., 2014, 2015). Meanwhile, the majority of cellular and transcriptomic work on the RSC has been carried out in mice (Brennan et al., 2020, 2021; Robles et al., 2020; Sullivan et al., 2023; Wang et al., 2023; Yamawaki et al., 2016, 2019a, 2019b; Yao et al., 2021, 2023), although there are examples of behavioral work in mice (Powell et al., 2020; Shi et al., 2024) and cellular work in rats (Kurotani et al., 2013; Q. Li et al., 2002; Yousuf et al., 2020). A systematic comparison of the cellular architecture of the RSC in both mice and rats would allow for a better understanding of retrosplenial circuit computation similarities and differences across the two species.

The RSC is divided into the granular (RSG) and dysgranular (RSD) subregions, which, while closely related, show significant functional and connectomic differences. For example, the RSG and RSD have been shown to play differential roles in contextual fear memory formation and recall (Pan et al., 2022; Tsai et al., 2022), and egocentric boundary vector cells are found primarily in RSD but not RSG (Alexander et al., 2020).

The two subregions are also differentially innervated by thalamic nuclei (Aggleton et al., 2021; Brennan et al., 2021; Lomi et al., 2021; Sripanidkulchai & Wyss, 1986). The RSG, in particular, is also remarkably distinct from other cortical regions in terms of its constituent cell types. RSG is home to a population of uniquely hyperexcitable pyramidal neurons, termed low-rheobase (LR) neurons, located exclusively in layers 2/3 (L2/3) (Brennan et al., 2020, 2021) and expressing the gene *Cxcl14* (Sullivan et al., 2023; Yao et al., 2021). Mouse RSG also contains a population of superficial L5 neurons marked by the gene *Scnn1a* and expressing an unusual mix of canonically intratelencephalic (IT) projecting and extratelencephalic (ET) projecting marker genes (Sullivan et al., 2023; Whitesell et al., 2021; Yao et al., 2021). It is currently unknown whether the cell types which distinguish RSG from RSD and neighboring cortical regions are conserved, lost, or amplified between mice and rats. Quantifying the degree to which the cell types of the mouse and rat RSG are conserved would shed light on whether the RSG’s transcriptomic distinctness has been selected for across evolutionary timescales.

Here we assess whether the distinct cell types of the mouse RSG are preserved in rats, if they are over- or underrepresented, and whether they are distinguished by the same patterns of synaptic, ion channel, and marker gene expression. We find that despite alterations in marker genes across species, two of the key unique cell types seen in the mouse RSG are not only preserved in rats, but also enhanced in terms of their proportional representation. These results highlight the evolutionary conservation of the unique RSG cell types, but also emphasize stark species-specific differences in marker genes for key cell types that need to be considered when making cell-type-specific transgenic lines of mice versus rats.

## MATERIALS AND METHODS

### Tissue dissections

All procedures and use of animals were approved by the University of Michigan Institutional Animal Care and Use Committee.

Animals were deeply anesthetized with isoflurane and swiftly decapitated, and the brains were moved into ice-cold ACSF (containing 126 mM NaCl, 1.25 mM NaH2PO4, 26 mM NaHCO3, 3 mM KCl, 10 mM dextrose, 1.20 mM CaCl2, and 1 mM MgSO4). Coronal slices of 500 um thickness for mice, and 800 um thickness for rats were obtained using Leica 1200 VT vibratome. Slices with anterior retrosplenial cortex (defined as AP -1.4 to -2.4 for mice and -2.2 to -4.0 for rat) were further selected. Microdissections of retrosplenial cortex from those slices was performed in the same but fresh ice-cold ACSF under Leica S9i stereo microscope. Retrosplenial cortex was dissected by making two straight, diagonal cuts running roughly from the highest point of the cingulum bundle to the pia at an angle following the reference atlas granular and dysgranular retrosplenial cortex border. One such cut was made on each hemisphere, and the tissue was then teased out along corpus callosum (CC), incorporating as little CC as possible. Tissue samples were snap frozen (one per animal) and held at -80 °C until nuclei isolation.

### Nuclei isolation

Retrosplenial cortex microdissections from up to two animals of the same group were combined into the same sample, to ensure sufficient cell count for sequencing. Each snap frozen sample was moved into a lysis buffer (EZ Prep Lysis Buffer with Protector RNAse inhibitor, Sigma) and manually homogenized in a glass dounce homogenizer. The sample was then filtered through a 30µm MACS strainer, cold centrifuged at 500 rcf per 5min and resuspended in the lysis buffer. This process was repeated twice more, except each time the pellet was resuspended in a wash buffer consisting of 5mM KCl, 12.5mM MgCl2, 10mM Tris Buffer pH 8.0, 1% BSA with RNAse inhibitor. After the last resuspension the sample was stained with propidium iodide to aid nuclei identification during flow sorting and was finally submitted to the University of Michigan Flow Cytometry Core for cell sorting.

### Library generation and read alignment

The 10x Genomics Chromium Single Cell 3’ Reagent Kit v3 was used to process the nuclei. Libraries were sequenced on a NovaSeq X system. Raw reads were aligned to the GRCm39 mouse reference genome (https://www.ncbi.nlm.nih.gov/datasets/genome/GCF_000001635.27/) or the mRatBN7.2 rat reference genome (de Jong et al., 2024) using Cell Ranger version 7.1 to generate counts matrices.

### Clustering

Clustering was performed independently on mouse and rat datasets using the Scanpy package (Wolf et al., 2018) in Python 3.10. First, putative doublets were detected using Scrublet (Wolock et al., 2019) and then removed. Cells with fewer than 750 total detected genes or fewer than 1000 total counts were additionally discarded. As our datasets were generated from single nuclei, little to no mitochondrial DNA is expected in uncontaminated nuclei, we removed cells with greater than 0.5% total mitochondrial gene counts. A round of initial clustering was then performed using the graph-based Leiden clustering algorithm to broadly classify cells as either neuronal or non-neuronal, for the purpose of removing clusters of non-neuronal cells from downstream analyses. Specifically, we utilized the Allen Brain Cell Atlas’s (ABCA) Whole Mouse Brain 10xv3 (WMB-10xv3) dataset (Yao et al., 2023) as a reference for mutual nearest neighbors (MNN) batch correction as described in (Haghverdi et al., 2018), followed by K nearest neighbors classification of each Leiden cluster. Only cells from the WMB-10xv3 dataset with the region label “RSP” were used as a reference. Each cell in our datasets was assigned a class from the WMB dataset based on the class of its 10 nearest neighbors in MNN-batch-corrected space. Each Leiden cluster was then assigned a class corresponding to that of the majority of its constituent cells. All cells falling within clusters corresponding to non-neuronal cell classes were discarded.

With only putative neurons remaining, we then performed an additional round of Leiden clustering. Again, using the WMB-10xv3 dataset with “RSP” cells selected, we assigned identities to each Leiden cluster according to the 10 nearest gaussian-weighted neighbors of each of its constituent cells in MNN-corrected space. The WMB-10x dataset contains four hierarchical levels of cluster classification, ranked “class,” “subclass,” “supertype,” and “cluster” from least to most finely detailed. A Leiden cluster was only assigned each of these classifications if at least 60% of its cells agreed on the classification, else it was marked as unclear. Clusters were then assigned a final identity according to the most detailed classification category on which at least 60% of its cells agreed, e.g. a Leiden cluster with a clear “supertype” designation but no clear “cluster” consensus would be assigned its identity according to its supertype. Each Leiden cluster was then merged with all other Leiden clusters with which it shared an assigned identity. GABAergic clusters (falling under either the “06 CTX-CGE GABA” or “07 CTX-MGE GABA” class designations) were merged according to their subclass designation. This resulted in a final total merged Leiden cluster count of 18 in mice and 18 in rats. We emphasize that this number is identical in the two datasets despite the clustering being performed entirely independently.

Mouse clusters with no consensus across any of the cluster, supertype, or subclass categories from the WMB-10x dataset were removed under the assumption that they corresponded to cell types not present in the reference dataset (in which we only selected cells from retrosplenial cortex), and therefore were likely only present in our transcriptomic data because of accidental inclusion of small amounts of neighboring brain regions during dissection. This exclusion was not applied to rat clusters, in order to avoid discarding potential rat cell types simply not found in mice. Instead, any rat clusters without a clear consensus were classified according to the cluster in our mouse dataset from which they derived at least 80% of their mutual nearest neighbors (see the MNN alignment section below), a threshold which all clusters met. We additionally removed clusters from both datasets corresponding to non-neuronal cell types that the initial round of clustering had failed to identify, as well as clusters with extremely low mean counts (less than 6000).

We then visualized the spatial distributions of all remaining merged Leiden clusters using the ABCA’s Whole Mouse Brain MERFISH (WMB-MERFISH) dataset. We found a very small, merged Leiden cluster in each of the mouse and rat datasets which corresponded to the “0087 L2/3 IT RSP Glut_1” supertype, which is uniquely localized to RSD in the WMB-MERFISH data. It was therefore removed on the basis that our microdissections had likely clipped a small amount of RSD, and the goal of this study is to characterize solely the cell types which are found in RSG. At the end of this process, 14 final clusters remained in both the mouse and rat datasets.

### MNN alignment of mouse and rat clusters

Identifying common cell types across multiple biological samples is a recurring problem in single-cell transcriptomics (Haghverdi et al., 2018; X. Li et al., 2020; Zhang et al., 2023). Relatively recently, a mutual-nearest-neighbors (MNN) based technique has been shown to excellently identify cells of the same type across samples where the inter-dataset variation, or “batch effect”, between the same cell types in two different samples is less than the intra-dataset variation between biological cell types (Haghverdi et al., 2018). We hypothesized that because mice and rats represent closely related species on evolutionary timescales, this principle would allow MNNs to identify analogous cell types between the two species.

We first obtained a subset of each dataset’s counts matrix for only the genes that mice and rats have in common. We then computed the log-normalized counts per million of these expression matrices and transformed them into cosine-normalized space. The K nearest neighbors (K=15) of each mouse cell in the rat dataset were computed, and vice versa. Any cross-species pairs of cells which were in each other’s K nearest neighborhood were counted as mutual nearest neighbors. These pairs of cells were then grouped by their respective clusters to generate the cross-species mutual nearest neighbors matrix.

### Co-clustering matrix

In order to further validate the fidelity of our MNN-based cross-species mapping, we generated a co-clustering matrix measuring the degree to which our mouse and rat clusters cluster together when the two datasets are combined. We first computed the top 1500 highly variable genes in each dataset using the scanpy function highly_variable_genes with the flavor=“seurat” option. We obtained the union of these two sets of genes, then subset each dataset for this union. In order to prevent species differences between analogous clusters from precluding legitimate co-clustering, we applied the MNN batch correction (K=20) from Haghverdi et al. (2018) to our rat dataset, using our mouse dataset as a reference.

We next generated 1000 random subsets of half of all cells from this combined, batch-corrected dataset. On each random subset, we first removed all genes which showed a high level of variation between species. We did this by Z-scoring genes between mouse and rat populations, removing all genes with a Z score above 0.5. We next computed the top 20 principal components (PCs) of each expression matrix, and using the same Z-scoring technique, removed PCs with a cross-species Z score above 0.25. We then used scanpy’s neighbors (n_neighbors=20, method=”gauss”) and leiden (resolution=1) functions to cluster each random subset.

Across all Leiden clusters in each subset, we counted the number of co-occurrences of cells from each original mouse cluster and rat cluster. These co-occurrences were added to a running total in the form of a two-dimensional matrix with mouse clusters on one axis and rat clusters on the other. For example, if a Leiden cluster contained 10 cells, 5 of which were mouse L2/3 LR cells, 4 of which were rat L2/3 LR cells, and 1 of which was a rat L6b CTX cell, the co-clustering matrix position at (Mouse L2/3 LR, Rat L2/3 LR) would increase by 5 x 4 = 20 and the position at (Mouse L2/3 LR, Rat L6b CTX) would increase by 5 x 1 = 5. The final co-clustering matrix consisted of each of these sums across all 1000 random subsets.

We analyzed three versions of the final co-clustering matrix: the raw matrix, one normalized by the sum of each row, and one normalized by the sum of each column. For each version, we then repeated the following process of assigning cluster mappings until each cluster was assigned a cross-species identity. First, we took the highest value in the matrix and assumed that its corresponding row (mouse cluster) and column (rat cluster) represented analogous cell types. We then removed that row and column from the matrix and returned to the first step, locating next the highest value in the matrix. All three methods assigned exactly the same cross-species cluster mapping as the MNN matrix had, thus validating our earlier results.

### Pearson correlation matrix

For each cluster in both species, we computed the mean expression vectors on a set of log-CPM-normalized highly variable genes. We then used these mean expression vectors to compute Pearson correlation coefficients on each cross-species pair of clusters. The set of highly variable genes was chosen by computing the top 1000 highly variable genes in both datasets using Scanpy’s highly_variable_genes function with the flavor="seurat" argument. We used the union of the two datasets’ highly variable genes, subsetting for only those genes present in both species.

Cross-species cluster mappings were obtained from the resulting correlation matrix using the same method as was used for the co-clustering matrix, described above.

### Cross-species similarity score

Cross-species similarity score for a given mouse-rat cluster pair on a specific group of genes was computed as the mean intracluster distance divided by the mean intercluster distance across all points in both clusters, as described in Chartrand et al. (2023). Data was cosine-normalized before computing this ratio. Where **D_a,b_** represents the mean of all individual distances between points in clusters **a** and **b**, and **D_a,a_** represents the mean distances between all points within cluster **a**, the cross-species similarity score for a mouse cluster **m** and rat cluster **r** is expressed as **(D_m,m_ + D_r,r_) / (2 * D_m,r_).**

The similarity matrix in **Supp. Fig. 1C** was computed on the same set of highly variable genes as described in the Pearson correlation matrix section, and cross-species cluster mappings were generated from the similarity matrix using the same technique described in the co-clustering matrix section.

### Experimental Design and Statistical Analyses

Tissue from a total of seven animals (three mice and four rats) was used to investigate retrosplenial transcriptomics. The following animals were used in this study: two male and two female five-month-old Sprague Dawley rats, and three four-month-old male C57BL6 background Pvalb-Cre (PV-IRES-Cre) mice. Only male mice were used, as previous transcriptomic studies of mouse cortex have already shown the absence of any major sex-specific cell type differences (Yao et al., 2021, 2023).

A Chi-square test was used to assess the significance of changes in cell type proportions between mice and rats. We used the proportions_chisquare function from the statsmodels 0.13.0 library (https://www.statsmodels.org/) in Python 3.10 to perform these tests. Effect sizes of cell type proportion differences were assessed using Cohen’s *h*, computed with Python code.

## RESULTS

### Ten excitatory cell types are found in the RSG, four of which uniquely define it

Previous work has identified several unique cell types within mouse RSG which are electrophysiologically, morphologically, and transcriptomically distinct from neighboring cortical regions (Brennan et al., 2020, 2021; Jedrasiak-Cape et al., 2024; Sullivan et al., 2023; Yao et al., 2021). These include a population of hyperexcitable L2/3 neurons known as LR cells (Brennan et al., 2020, 2021) which have been proposed to correspond to a transcriptomic *Cxcl14+* population of L2/3 RSG cells (Jedrasiak-Cape et al., 2024; Sullivan et al., 2023); a L5a population uniquely labeled by the gene *Scnn1a* (Sullivan et al., 2023; Yao et al., 2021); a deep L5 population uniquely marked by the gene *C1ql2* (Sullivan et al., 2023; Yao et al., 2021); and others. Our aims were twofold: To transcriptomically characterize the full complement of cell types found in the mouse RSG, and to determine the degree to which these cell types are evolutionarily conserved between mice and rats, groups which diverged more than 10 million years ago (Adkins et al., 2001; Nei et al., 2001; Springer et al., 2003).

We first utilized the Allen Institute’s Whole Mouse Brain Multiplexed Error-Robust Fluorescence in situ Hybridization (WMB-MERFISH) dataset (Yao et al., 2023) to visualize cell populations which comprise the mouse RSG **(Fig. 1A-B).** By examining WMB-MERFISH clusters which are found in RSG, we identified ten main excitatory cell types. We assigned names to these cell types according to a combination of their spatial identities and existing cell type annotations from the WMB-MERFISH dataset. In the case of a dense L2/3 *Cxcl14*-positive cell type which likely corresponds to LR cells, we additionally deferred to existing nomenclature for RSG cell types. This cell type, specifically the “0088 L2/3 IT RSP Glut_2” supertype in the WMB-MERFISH dataset, was one of two main glutamatergic cell types we identified in L2/3 of the examined MERFISH slices **(Fig. 1C)**. Its uniquely high density and spatial restriction to L2/3 RSG makes it by far the most compelling candidate for the transcriptomic correlate of electrophysiological LR neurons (Brennan et al., 2020, 2021; Sullivan et al., 2023; Jedrasiak-Cape et al., 2024), and we thus termed it “L2/3 LR”. The other, far sparser L2/3 cell type corresponds to a supertype named “0030 L2/3 IT CTX Glut_2,” which we shortened to “L2/3 IT CTX”. We next examined L5, finding four main cell types **(Fig. 1D)**. Superficial L5 is dominated by the supertype “0089 L4 RSP-ACA Glut_1,” which is found solely within RSG and which we therefore termed “L5a RSG.” Cross-referencing this supertype in the WMB-10xv3 dataset, we found that among RSC neurons it uniquely expresses the gene *Scnn1a*, a known marker gene for superficial L5 cells in RSG (Sullivan et al., 2023; Whitesell et al., 2021; Yao et al., 2021). Slightly deeper than the L5a RSG cell type lies the cluster “0060 L5 IT CTX Glut_2.” This cluster is found along much of medial neocortex (mCTX) including retrosplenial, anterior cingulate, and secondary motor cortices and we thus termed it “L5 IT mCTX.” In deep L5, we next identified two cell types, one a putatively near-projecting (NP) supertype named “0122 L5 NP CTX Glut_1” found almost exclusively in RSG which we termed “NP RSG”, and one a putatively extratelencephalic (ET) projecting cluster named “0381 L5 ET CTX Glut_6”, which we termed “L5 ET RSC” for its near-complete spatial restriction to RSC as a whole. Looking finally at L6, we again found four main resident cell types **(Fig. 1E)**. By far the most numerous of the L6 cell types are two putative corticothalamic (CT) clusters, one of which, “0450 L6 CT CTX Glut_4,” is spatially restricted to RSG, and which we thus termed “CT RSG”. The other, “0445 L6 CT CTX Glut_3,” is found along much the same range as the L5 IT mCTX cell type, and we thus refer to it as “CT mCTX”. The final two cell types are very sparse. One corresponds to the cluster “0041 L6 IT CTX Glut_2”, which shows little spatial restriction other than being confined to L6, and is here shortened to “L6 IT CTX”. The other is the subclass “029 L6b CTX Glut”, which encompasses all neocortical L6b neurons.

**Figure 1.**
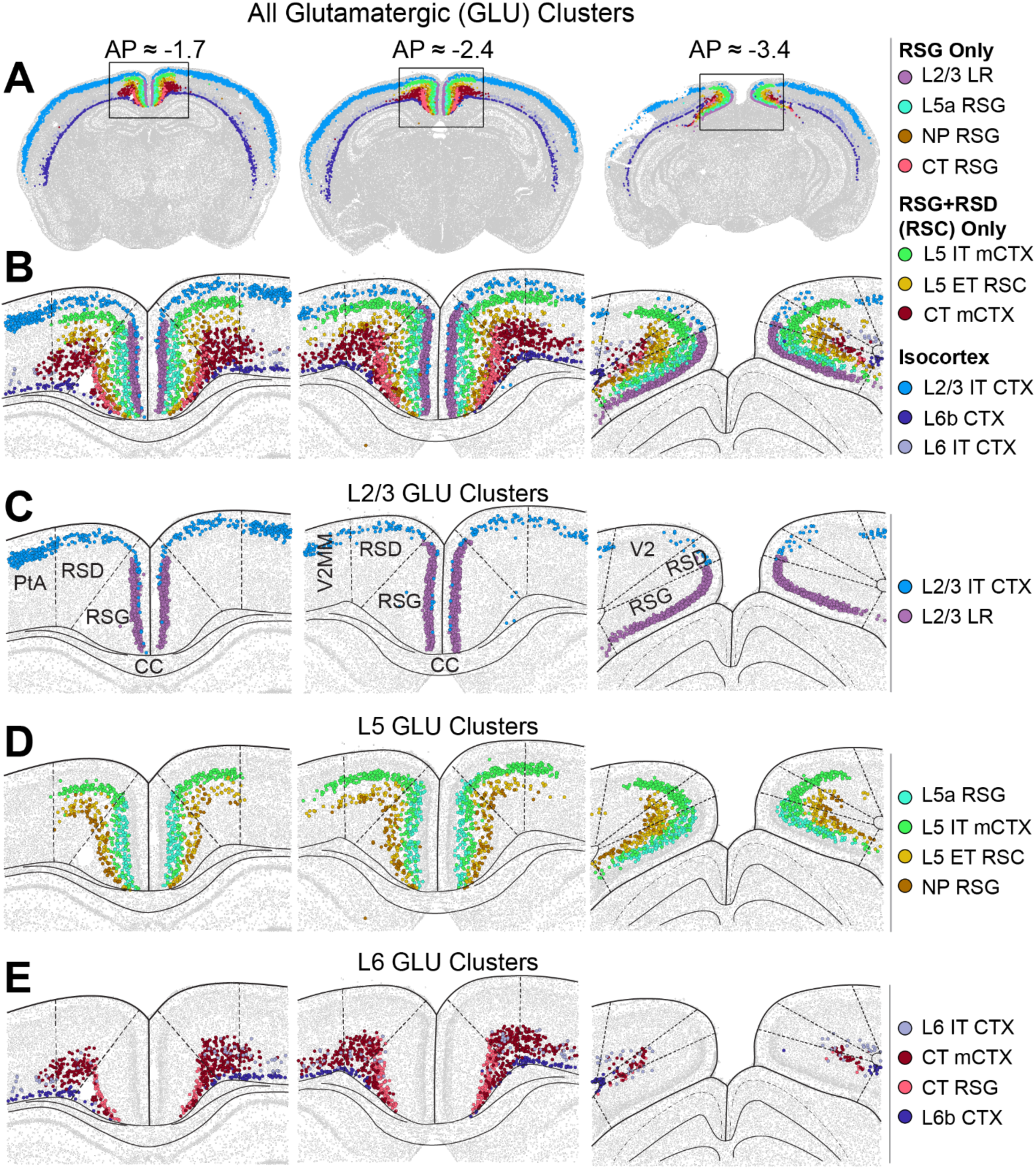
MERFISH distributions of excitatory cell types within RSG. (A) Selected coronal slices from the Allen Brain Cell Atlas MERFISH dataset showing the distributions of all 10 identified excitatory (glutamatergic) cell types in mice. (B) Zoomed in views of retrosplenial cortex, showing the same slices and cell types as (A). (C to E) The same as (B), but with only L2/3 (C), L5 (D), or L6 I clusters shown for clarity. Note that expression of 4 of the 10 excitatory subtypes is nearly entirely restricted to only the RSG (L2/3 LR, L5a RSG, NP RSG, CT RSG).

### The same cell types are identified in mouse and rat RSG

We next sought to determine the degree to which these cell types are evolutionarily conserved between mice and rats. To do this we utilized two scRNA-seq datasets **(Fig. 2A)**: one newly generated from the RSG of rats, and one from the RSG of mice which we previously used in Jedrasiak-Cape et al. (2024). We performed quality control and unsupervised clustering independently on each dataset. After the full quality control and clustering process, 14 mouse and 14 rat clusters remained (See methods). Of these clusters, 10 were putatively excitatory and 4 were inhibitory in both species. We first sought to understand the identities of the clusters from our mouse dataset. Briefly, we utilized a mutual nearest neighbors (MNN) based batch-correction method (Haghverdi et al., 2018) to align our mouse data with the Allen Institute’s Whole Mouse Brain 10xv3 (WMB-10xv3) scRNA-seq dataset (Yao et al., 2023) and subsequently obtain identities for our mouse clusters (see methods). Crucially, the WMB-10xv3 and WMB-MERFISH datasets have already been clustered integratively, and it is therefore possible to confidently obtain information about the same cell population from both references. We were thus able to compare our own transcriptomic cell types with those we had previously identified in the WMB-MERFISH dataset. Our 10 excitatory mouse clusters matched one-to-one onto each of the 10 previously identified WMB-MERFISH cell types **(Fig. 2B)**. We thus assigned names to each of our excitatory mouse clusters based on the names we had given to their corresponding MERFISH cell type. Three of our four inhibitory clusters also cleanly mapped onto single subclass categories from the WMB dataset. We termed these clusters Pvalb, Sst, and Lamp5, as they respectively mapped onto the WMB dataset “052 Pvalb”, “053 Sst”, and “049 Lamp5” subclasses. However, one of our inhibitory clusters contained substantial numbers of cells from both the “046 Vip Gaba” and “047 Sncg Gaba” subclasses, and we therefore named this cluster Vip/Sncg.

**Figure 2.**
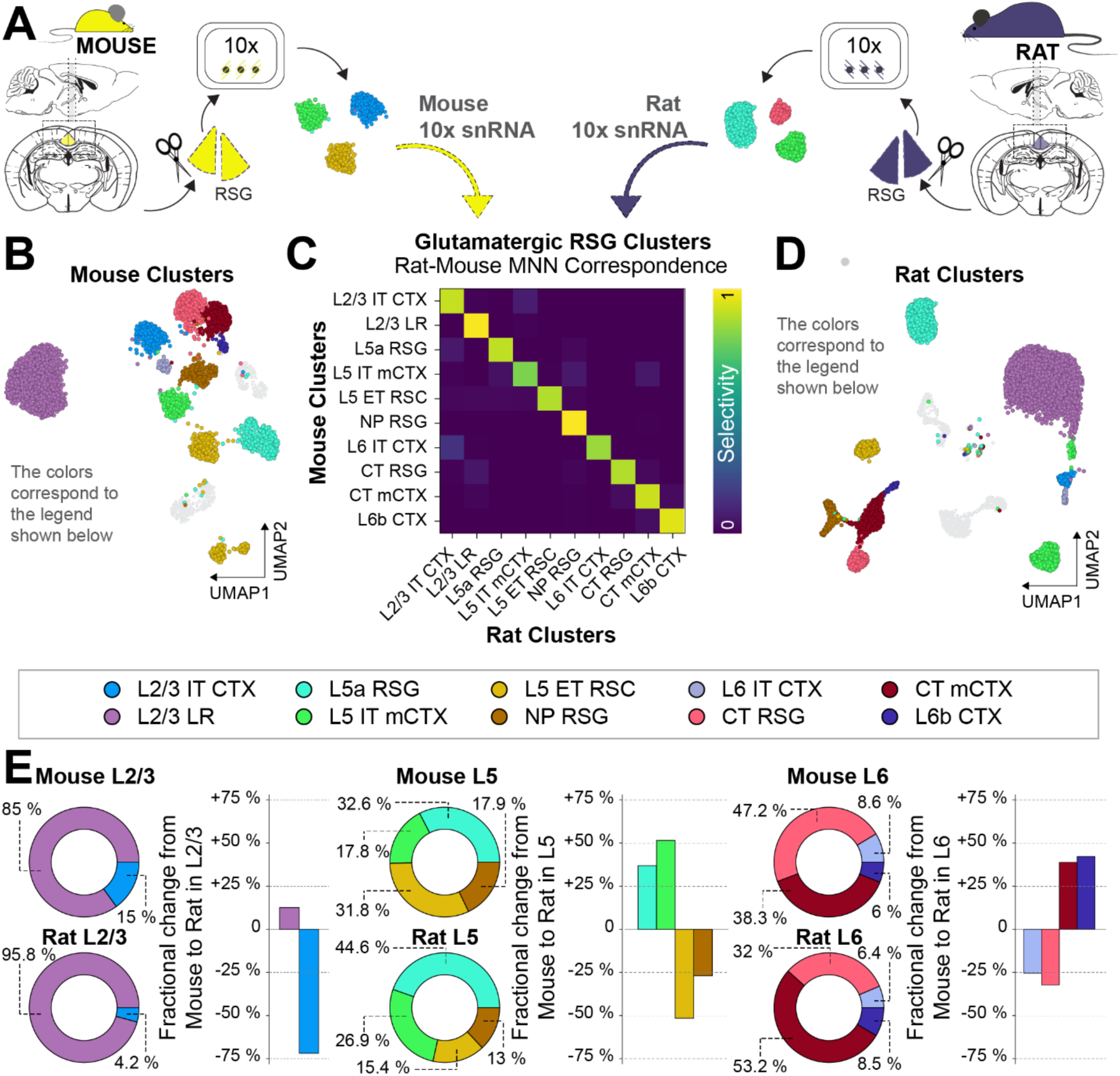
Clustering and alignment of mouse and rat RSG excitatory neurons. (A) Schematic showing the independent dissection and processing of mouse and rat snRNA-seq samples. (B) UMAP projection of mouse cells, colored by excitatory (glutamatergic) cluster identity. Inhibitory clusters are greyed out. (C) Mutual nearest neighbor correspondence matrix of mouse and rat excitatory clusters. For a given mouse cluster M and rat cluster R, the value shown is the proportion of R’s mutual nearest neighbors in the mouse dataset which are from M. (D) UMAP projection of rat cells, colored by cluster identity. Inhibitory clusters are greyed out. I Proportions of cell types within specific layers in mice versus rats (pie charts), and fractional change of within-layer cell type proportions in rats compared to mice (bar charts). For a given cell type with proportion M in mice and R in rats, fractional change percentage is computed as ((R / M) – 1) * 100%. Significance of proportion changes of each cell type within each layer were assessed using a Chi-square test, and effect sizes were computed as nondirectional Cohen’s *h*. P values and effect sizes are as follows: L2/3 IT CTX and L2/3 LR (the only two groups in L2/3), p = 1.8e-51, *h* = 0.40; L5a RSG, p = 4.5e-11, *h* = 0.24; L5 IT mCTX, p = 4.7e-9, *h* = 0.21; L5 ET RSC, p = 1.26e-25, *h* = 0.37; NP RSG, p = 9.6e-5, *h* = 0.14; L6 IT, p = 0.085 *h* = 0.083; CT RSG, p = 1.4e-10, *h* = 0.31; CT mCTX, p = 9.5e-10, *h* = 0.30; L6b CTX, p = 0.049, *h* = 0.098.

Having assigned identities to our 14 mouse clusters, we next characterized the degree to which our 14 rat clusters represented analogous, evolutionarily conserved cell types between the two species. We focused initially on determining the level of correspondence of the 10 excitatory clusters in each dataset. To do this, we again utilized a mutual nearest neighbors-based technique (see methods). Grouping MNNs by cluster in each dataset, we observed a clear one-to-one correspondence between mouse and rat clusters **(Fig. 2C)**, confirming that the whole suite of 10 glutamatergic RSG cell types is conserved. In order to further validate this result in additional ways, we employed three additional methods of mapping cluster identity across species. We generated a co-clustering matrix of mouse and rat clusters by repeatedly clustering random subsets of both datasets together **(Supp. Fig. 1A)**, we computed the Pearson correlation coefficient on mean expression vectors between each cross-species cluster pair **(Supp. Fig. 1B)**, and finally we computed a cluster similarity score as described in (Chartrand et al., 2023), equivalent to the ratio of the mean intracluster distance to the mean intercluster distance, on each pair of clusters **(Supp. Fig. 1C)**. Each method agreed fully with our initial MNN-based results (see methods). We then assigned names to each rat cluster **(Fig. 2D)** based on the mouse cluster with which it consistently aligned in all four of the mapping methods described.

With a common suite of excitatory cell types established between mouse and rat RSG, we next inspected the composition of the RSG by cortical layer **(Fig. 2E)**. Within L2/3 mouse clusters, we found that the proportion of L2/3 LR to L2/3 IT CTX cells, at 85% to 15% respectively, is remarkably in line with previously reported proportions of electrophysiological LR to RS populations within mouse L2/3 *in vitro* (Brennan et al., 2020). Surprisingly, however, this ratio is significantly altered (p = 1.8e-51, Chi-square test) in our rat dataset, at 95% to 5%, representing a substantial increase in L2/3 LR cells and a concomitant decrease in putative non-LR L2/3 cells. Within rat L2 RSG, slice electrophysiology has identified 94% of neurons as exhibiting a late-spiking phenotype (Kurotani et al., 2013), a feature known to be characteristic of LR cells in mice (Brennan et al., 2020). This suggests that our transcriptomic data precisely reflects real biological proportions of cell types in RSG. Compared to mouse L5, in rats we found substantial and significant increases in the proportions of the more superficial cell types, L5a RSG (35% increase, p = 4.5e-11) and L5 IT mCTX (50% increase, p = 4.7e-9), accompanied by proportional decreases in deeper cell types, L5 ET RSC (49% decrease, p = 1.26e-25) and NP RSG (28% decrease, p = 9.6e-5). L6 saw a decrease in the proportion of CT RSG (32% decrease, p = 1.4e-10), and an increase in CT mCTX (39% increase, p = 9.45e-10) with non-significant changes in the L6 IT CTX (25% decrease, p = 0.085) and L6b CTX (42% increase, p = 0.049) clusters at the chosen significance threshold of 0.01.

We next focused on the four inhibitory clusters in both datasets. In mice, these clusters consist of the MGE-derived Pvalb and Sst classes of interneurons, and the CGE-derived Vip/Sncg and Lamp5 classes **(Fig. 3A-B)**, so named for genes which they are known to reliably and selectively express (Gouwens et al., 2020; Lim et al., 2018; Tasic et al., 2018). As all four of these groups are evolutionarily conserved between mice and humans (Hodge et al., 2019; B. Kim et al., 2023), we predicted that rats would have a 1:1 correspondence in their inhibitory clusters, just as with the excitatory clusters. Indeed, upon computing the same MNN-based correspondence, we observed a clean 1:1 mapping of these inhibitory cell types between mice and rats **(Fig. 3D).** The proportions of inhibitory subtypes showed some variation between the two species, however the only statistically significant change at our chosen significance threshold of 0.01 was an increase in the proportion of the Pvalb cluster (22% increase; p = 0.0045, Chi-square test). The proportions of the Sst and Vip/Sncg clusters were slightly decreased in rats, although statistically insignificantly (Sst, 18% decrease, p = 0.05; Vip/Sncg, 13% decrease, p = 0.16). The proportion of the Lamp5 subtype remained relatively unchanged (1.2% increase, p = 0.93). We next examined the overall relative proportions of inhibitory and excitatory neurons in mouse and rat RSG. We observed a significant increase in the share of inhibitory neurons (18%, p = 0.0015), a finding with significant implications for circuit dynamics in the two species (Ahmed & Mehta, 2009).

**Figure 3.**
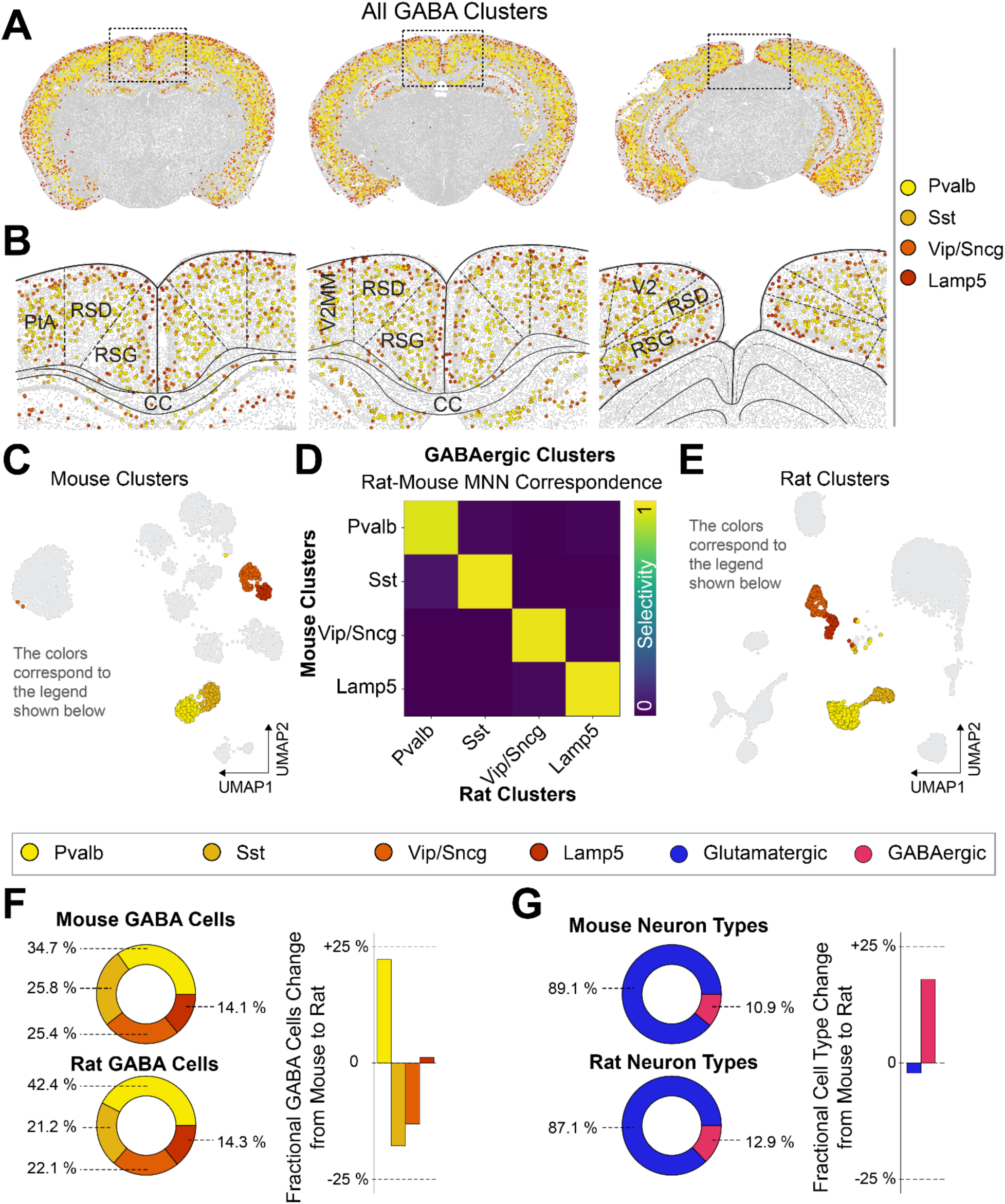
Overview of RSG inhibitory clusters. (A) Selected coronal slices from the Allen Brain Cell Atlas MERFISH dataset showing the distributions of 4 major inhibitory (GABAergic) subtypes in mice. (B) Zoomed in views of retrosplenial cortex, showing the same slices and cell types as (A). (C) UMAP projection of mouse cells, colored by cluster identity. Glutamatergic clusters are greyed out. (D) Mutual nearest neighbor correspondence matrix of mouse and rat inhibitory clusters. (E) UMAP projection of rat cells, colored by cluster identity. Glutamatergic clusters are greyed out. (F) Proportions of classes of inhibitory neurons in both mice and rats (pie charts), and fractional change in cell type proportions in rats as compared to mice (bar charts). Significance of proportion changes of each cell type within each layer were assessed using a Chi-square test, and effect sizes were computed as nondirectional Cohen’s *h*. p values and effect sizes are as follows: Pvalb, p = 0.0045, *h* = 0.16; Sst, p = 0.049, *h* = 0.11; Vip/Sncg, p = 0.16, *h* = 0.08; Lamp5, p = 0.93; *h =* 0.0079. (G) Proportions of total inhibitory and excitatory populations in both mice and rats (pie charts), and fractional changes in total inhibitory and excitatory population proportions in rats as compared to mice. Significance was assessed the same way as in (F): p = 0.0015, *h* = 0.061.

### Key marker genes are not reliably conserved across species

Marker genes are defined as genes which exhibit a relatively high degree of anatomical or cell type specificity in their expression pattern. Identification of marker genes is essential for performing cell-type-specific manipulations using genetic tools such as the Cre/lox system combined with knock-in optogenetic reporters, a nearly ubiquitous technique in the study of model organisms. In mice and rats, two closely related model organisms, engineered Cre lines for certain evolutionarily conserved marker genes have been used to reliably target the same classes of cells across both species (Witten et al., 2011). We explored the degree to which known marker genes for mouse RSG cell types are conserved in rats, as this information is a crucial pre-requisite to selective targeting of specific cell types in both species.

In mice, the genes *Cxcl14* and *Scnn1a* have previously been identified as a marker genes of the RSG-specific L2/3 LR and L5a RSG cell types, respectively (Sullivan et al., 2023; Yao et al., 2021). We confirmed the same pattern of expression in our mouse dataset, but surprisingly, the expression of these genes differed substantially in rats. Specifically, expression of the *Scnn1a* marker gene was completely absent in the rat L5a RSG cell type, despite strong labeling of the same cluster in mice. **(Fig. 4A).** *Cxcl14*, while still relatively selectively expressed in the L2/3 LR cluster in both species, was present in far lower levels in rats **(Fig. 4B)**. Given the failure of *Scnn1a* to label L5a RSG cells in rats, and the significantly reduced expression of *Cxcl14* in rat L2/3 LR cells, we examined the rat dataset for other candidate marker genes for these clusters **(Fig. 4B).** In LR cells, we identified the marker gene *Shisa8,* which to our knowledge has not previously been known to mark LR cells. This gene is expressed at similar levels in both mice and rats and represents a compelling candidate for future genetic manipulations of LR cells in both species. For the L5a RSG cluster, we examined two genes previously shown to be enriched in this cell type in mice, *Gpr6* (Sullivan et al., 2023) and *Hsd11b1* (Yao et al., 2021). We found that the expression pattern of these genes was extremely well conserved between species, and we thus propose both genes as possible candidates for genetic targeting of L5a RSG cells in rats.

**Figure 4.**
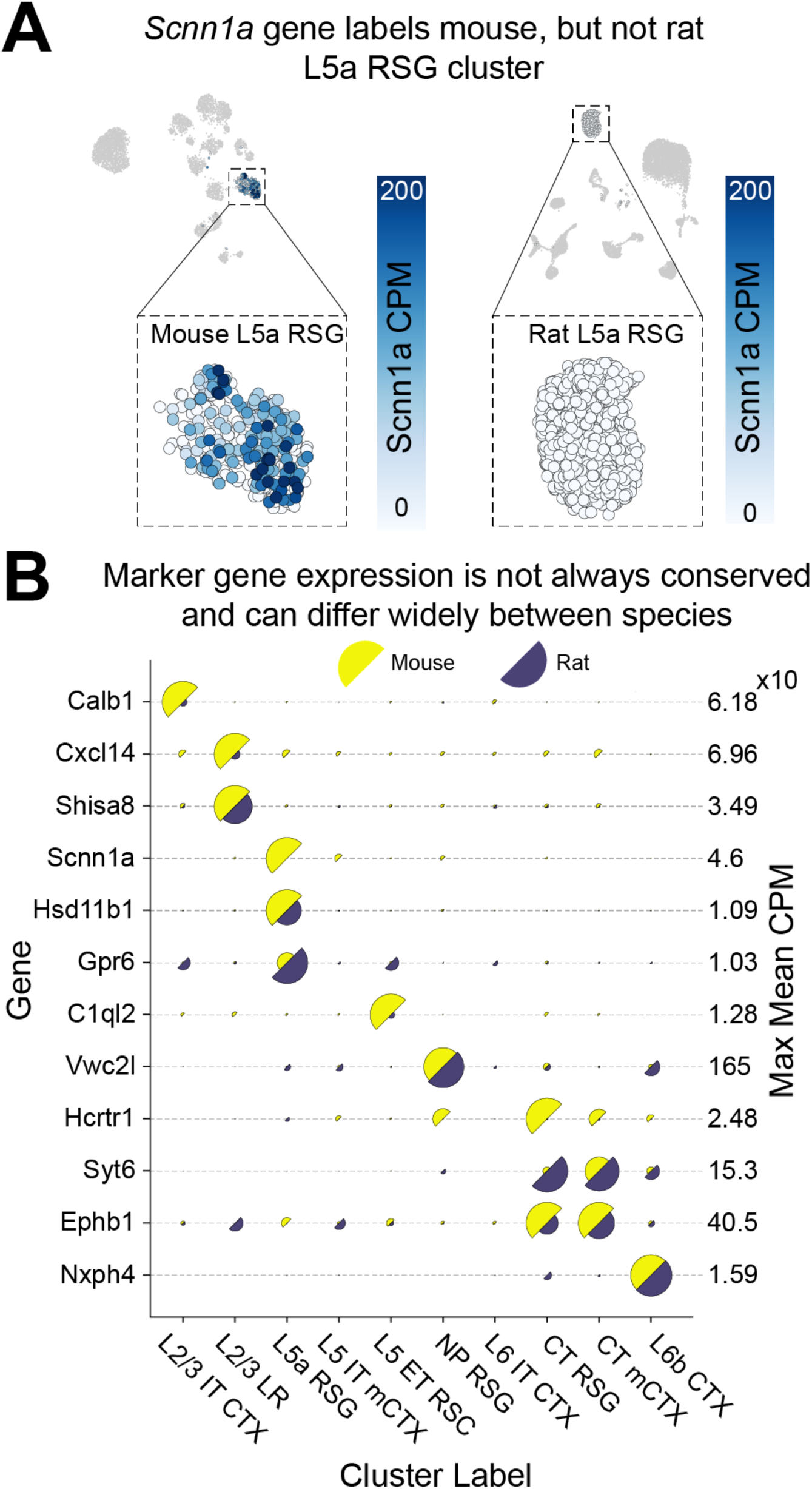
Marker genes for specific RSG cell types are not consistently conserved across species. (A) UMAP projections of mouse (left) and rat (right) cells, with the L5a RSG cluster highlighted. L5a RSG cells are color-coded according to expression of *Scnn1a*, a known marker gene for this cluster in mice. Note the complete absence of this mouse marker gene in rat L5a RSG cells. (B) Half-circle plots of selected marker gene expression in excitatory clusters. Semicircle sizes are normalized by the maximum value in each row (corresponding to the maximum value for a given gene). The far-right column of the panel denotes the value represented by the largest semicircle in each row.

We additionally examined the expression of known inhibitory marker genes in GABAergic clusters **(Supp. Fig. 2)**, finding that canonical marker genes remain extremely well conserved between mice and rats. This is true despite the fact that overall expression of the *Pvalb* gene within the Pvalb cell type is much lower in rats than in mice: The proportion of cells in this cluster expressing *Pvalb* was altered only slightly, at 91% expression in rats and 81% in mice.

### Transcriptomic profile of neurotransmitter receptors remains largely conserved, with some key differences

The expression of neurotransmitter receptors determines how neurons respond to chemical synaptic input. We examined the expression of glutamate, serotonin, and muscarinic acetylcholine receptors across mouse and rat datasets in order to understand the degree to which RSG circuit dynamics may be preserved.

Notably, the L2/3 LR and L5a RSG clusters were found to almost entirely lack expression of the gene *Gria1*, which encodes the GluA1 AMPA receptor subunit **(Fig. 5A)**. AMPA receptors commonly assemble into heterotetramers composed of combinations of at least two of four possible subunits, GluA1-4 *(Gria1-4)* (Traynelis et al., 2010). While it is unclear what functional purpose the lack of GluA1 serves, it is however known that presence of the GluA2 subunit confers calcium (Ca2+) impermeability in the fully assembled tetramer (Cull-Candy & Farrant, 2021). Given the extremely high intrinsic excitability of LR cells, it is possible that the lack of GluA1 has the effect of avoiding calcium-dependent excitotoxicity by enabling a higher proportion of AMPA receptors to assemble with the GluA2 subunit.

**Figure 5.**
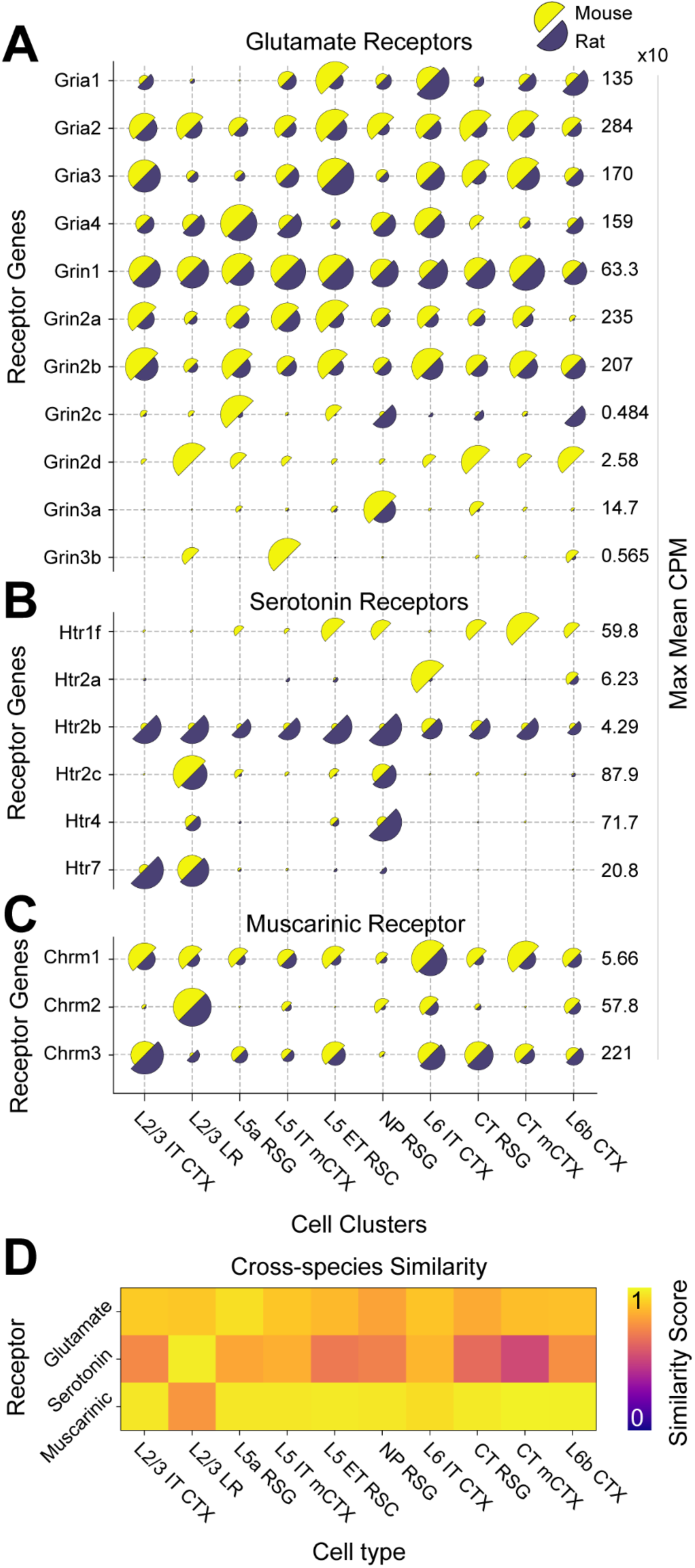
Expression of neurotransmitter receptors remains largely conserved across mouse and rat RSG excitatory clusters, with key differences. (A to C) Half-circle plots of selected neurotransmitter receptor categories in excitatory clusters. Within each row, the radius of each semicircle is proportional to mean CPM of that row’s gene. The far-right column of each plot contains the value represented by the largest semicircle in each row. (D) Cross-species similarity scores (see methods) for each cluster across all neurotransmitter receptor categories.

We noted several other key features of neurotransmitter receptor expression in LR cells which were conserved across species. In both mouse and rat RSG, LR cells express the highest levels of the serotonin receptor *Htr2c*, and also express much higher levels of serotonin receptors *Htr4* and *Htr7* than most other cell types **(Fig. 5B).** LR cells are also characterized in both species by their uniquely high level of muscarinic acetylcholine receptor *Chrm2* **(Fig. 5C)**. All of this indicates that within the neurotransmitter systems examined here, many key features which distinguish LRs from other cell types remain remarkably well conserved. We may therefore reasonably expect neuromodulation of LR cells to behave similarly in mice and rats, supporting the idea of an evolutionarily stable computational role for this cell type across the two species.

Several surprising trends emerged upon examining the expression of serotonin receptors across other cell types. Most notably, expression of the *Htr1f* gene, which in our mouse data is highly expressed in all deep L5 and L6 clusters except L6 IT CTX, is entirely lacking in our rat dataset. There appears to be little existing literature on the function of the 5-HT_1F_ receptor in neurons, and it is thus difficult to address how the lack of this receptor in rat RSG could affect circuit dynamics. The *Htr2a* gene, which encodes the 5-HT_2A_ receptor and is commonly studied within the field of psychedelic research (Ekins et al., 2023; Johnson et al., 2019), is almost completely lacking from all cell types in both mouse and rat RSG. The mouse L6 IT cluster expresses by far the highest level of this receptor, but interestingly almost completely lacks it in rats. Meanwhile, expression of *Htr2b* shows an increase in rats relative to mice across all cell types and *Htr2c* remains largely the same, expressed mostly in the L2/3 LR and NP RSG clusters.

We additionally compared neurotransmitter receptor gene expression in inhibitory neurons **(Supp. Fig. 3)**, finding that, as with excitatory neurons, general expression patterns remained conserved while specific genes such as *Htr1f* showed surprising cross-species differences.

### Ion channel expression offers insights into electrophysiological and computational properties

The electrophysiological properties of neurons are determined by ion channels embedded in the cell membrane. These include voltage-gated sodium (Na+) and potassium (K+) channels which determine the properties of a neuron’s action potentials, as well as non-voltage-gated K+ and Na+ leak channels primarily responsible for determining the resting membrane potential. We examined the expression of genes encoding these channels, finding broad expression patterns that were consistent across mice and rats, as well as specific genes that differed notably.

LR cells exhibit a late-spiking property, a feature that, when compared to neighboring excitatory cell types, is more pronounced in rats than in mice (Brennan et al., 2020; Kurotani et al., 2013). In rats, this property has been shown to depend on the Kv1.1 *(Kcna1)*, Kv1.4 *(Kcna4)*, and Kv4.3 *(Kcnd3)* voltage-gated potassium channels (Kurotani et al., 2013). While each of these genes is expressed in the L2/3 LR cluster in both mice and rats, our dataset shows much higher expression of these genes in LR cells in rats than in mice **(Fig. 6A).** This is true for percent expression of these genes in addition to overall mean expression levels; for example, only 35% of mouse LR cells express *Kcna4* in our dataset, while 94% of rat LR cells do. This substantial increase in the expression of relevant potassium channels offers a potential explanation for why the late-spiking property may be much more pronounced in rats.

**Figure 6.**
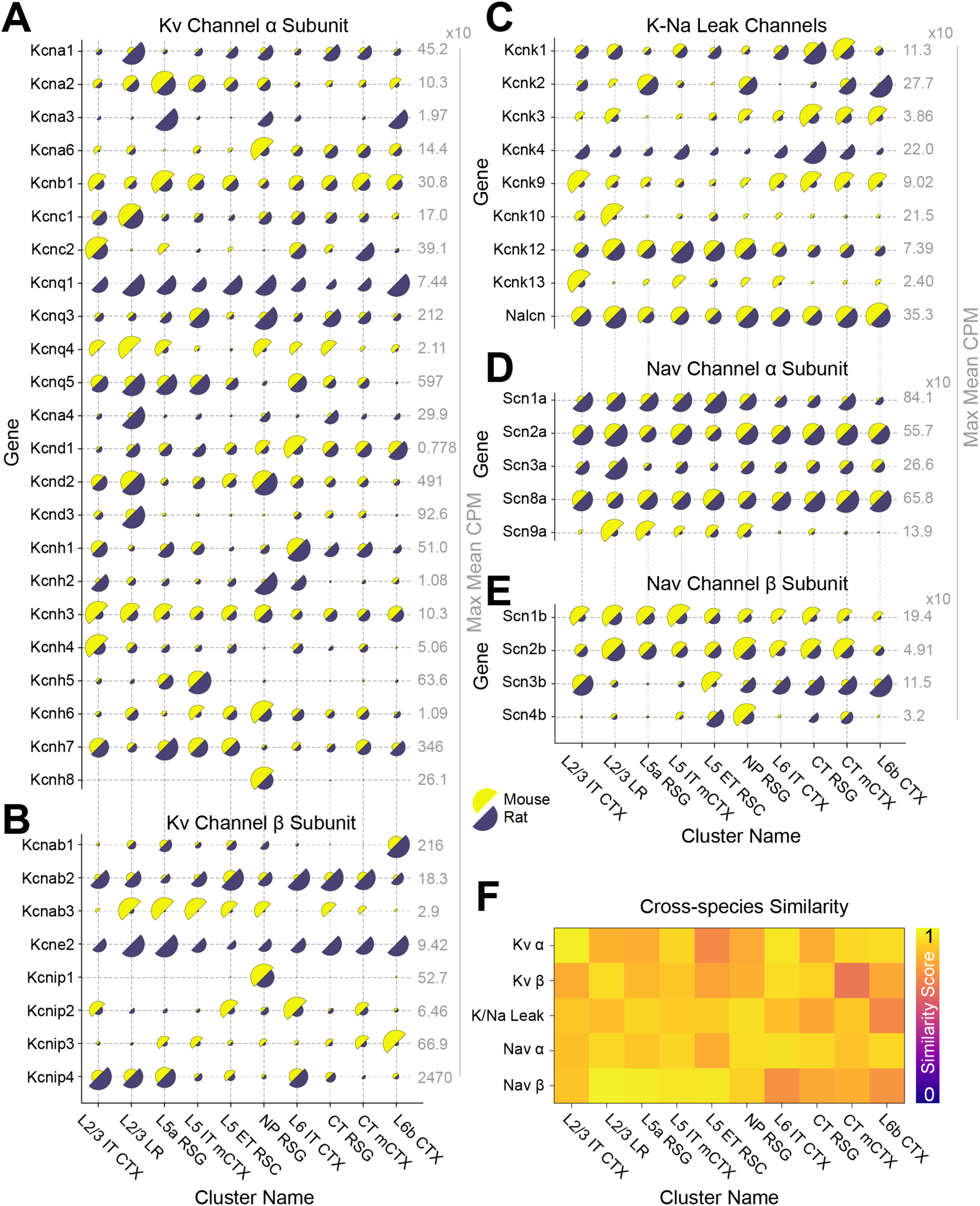
Expression of sodium and potassium ion channels across mouse and rat RSG clusters. (A to E) Half-circle plots of selected ion channel categories in excitatory clusters. Within each row, the radius of each semicircle is proportional to mean CPM of that row’s gene. The far right column of each plot contains the value represented by the largest semicircle in each row. (F) Cross-species similarity scores (see methods) for each cluster across all ion channel categories.

A few specific genes showed very different expression across all or most cell types between mice and rats. For example, the voltage-gated K+ channel β subunit *Kncab3* is nearly ubiquitously expressed at relatively low levels in the mouse dataset, but is almost entirely missing from rats. The reverse is true for *Kcne2*, a β subunit responsible for a host of strong modulatory effects on voltage-dependent potassium channels (Abbott, 2012). Among K+ leak channels, *Kcnk4* is missing from mice but expressed ubiquitously in rats, and *Kcnk9* is found in all mouse clusters but sharply underexpressed in rats. The implications of these differences in K+ channel expression for mouse and rat RSG cell physiology remain to be examined, but may result in altered resting membrane potentials or action potential dynamics across the two species.

We examined expression of these same genes for inhibitory neurons, again finding broad similarities across mice and rats **(Supp. Fig. 4).** Several differences observed in excitatory cell types, such as rat-specific expression of *Kcne2* and mouse-specific expression of *Kcnk4*, were also observed in inhibitory cell types. While our dataset only encompasses the RSG, the fact that these genes exhibit such stark changes in expression across all cell types, including cell types which are also found elsewhere in the cortex, may be indicative of wider differences in expression patterns not specific to RSG.

## DISCUSSION

Here we show that the RSG in mice and rats is defined by a suite of evolutionarily conserved neuronal cell types. When it comes to inhibitory neuronal subtypes, this is perhaps an expected outcome as the GABAergic Pvalb, Sst, Vip/Sncg, and Lamp5 interneuron classes are found throughout the cortex (Lim et al., 2018; Tasic et al., 2018; Yao et al., 2021, 2023) and have each previously been identified in humans (Hodge et al., 2019; B. Kim et al., 2023), a species far more evolutionarily removed from mice than rats.

The mouse RSG’s excitatory (glutamatergic) cell types, however, are remarkably unique in comparison to even neighboring cortical regions (Brennan et al., 2020, 2021; Sullivan et al., 2023; Yao et al., 2021). The premise of our study was based on the reasoning that if these unique cell types in mice are truly of significance in achieving the RSG’s computational and behavioral functions, then they should be robustly preserved or enhanced in rats. These unique mouse excitatory cell types include the defining cell type of the RSG (L2/3 LR cells; (Brennan et al., 2020), as well as the L5a RSG population which in mice is selectively labeled by the gene *Scnn1a*. Both of these cell types are only found in the RSG and not in the adjacent RSD (Sullivan et al., 2023; Whitesell et al., 2021; Yao et al., 2021). We found that not only are both these cell types preserved in rat RSG, but they are enhanced in terms of the fraction of total excitatory cells they represent in their respective layers. This suggests that these cells are likely serving evolutionarily important computational and behavioral functions.

### Relevance for future study of disease state transcriptomes

In humans, the RSC is often studied in the context of Alzheimer’s disease (AD), as it is one of the earliest regions of the brain heavily affected by the condition (Choo et al., 2010; Minoshima et al., 1997). Even in preclinical rodent models of AD such as the 5xFAD mouse line, the RSC has been shown to be heavily impacted early on in disease progression (Gouwens et al., 2020). By transcriptomically characterizing the suite of cell types one can expect to find within the RSG, our work offers a framework through which to study the cell-type-specific transcriptomic effects of relevant disease models.

### A conserved cellular architecture underlies RSG computations in mice and rats

Similar cellular architectures should be expected to generate similar functional rhythmic circuit outputs across species. Cellular architecture places important constraints on brain-state-dependent rhythmic activation, and a circuit’s rhythm generation therefore serves as a useful computational output that can be compared across species. So, do mouse and rat RSG circuits generate similar brain rhythms in similar brain states, as would be expected from the existence of similar cell types across the two species? During both awake behaving states and REM sleep, theta rhythms are seen across most cortical regions, including the RSG (Alexander et al., 2020; Ghosh et al., 2022; Koike et al., 2017). What sets the RSG apart in terms of these theta states is the existence of a remarkably fast 110-160 Hz theta-coupled rhythm (splines) that is almost perfectly anti-phase across the left and right hemispheres, representing a unique form interhemispheric rhythmic communication that is strongest in the RSG (Ghosh et al., 2022). These rhythms are named after mechanical splines, the teeth on mechanical gears that are also anti-phase when they interlock across two interacting gears. Gamma rhythms (30-80 Hz) are also seen in the same theta cycles as splines, but on the rising phase of theta. Importantly, unlike the unique spline rhythms that are anti-phase across the left and right brain, these RSG gamma rhythms are almost perfectly in-phase across the two hemispheres. Our previous work has shown that the balance between anti-phase splines and in-phase gamma rhythms changes across brain states in both rats and mice in quantitatively similar ways (Ghosh et al., 2022; their Figure 4). That the nuanced brain-state-dependent features of this RSG-specific oscillation (splines) have been preserved so well between mice and rats strongly suggests preservation of much of the underlying circuitry, and therefore the constituent cell types. The broad similarities of ion channel and neurotransmitter gene expression patterns between mouse and rat RSG cell types further support the conclusion that the same circuitry underlies the RSG’s computational capacities in the two species, and also that the RSG performs relatively similar brain-state-dependent computations across mice and rats.

### Implications for Cre-driver line development

The ability to selectively manipulate cell types is an extremely powerful tool for investigating neural circuit function. Our work confirms several marker genes which consistently define the same cell types across species, but also identifies some which crucially do not. The data here suggests that the naïve development of *Cre* lines in rats based only on mouse cell-type markers could lead to unexpected results without prior evidence that a driving gene displays the same expression pattern in rats and mice. For example, if one were to develop an *Scnn1a*-Cre line in rats for the intended purpose of manipulating L5a RSG neurons, the evidence presented here suggests that the effort would be entirely unsuccessful. We have thus presented alternative marker genes to those already identified for both the L2/3 LR (marked by *Shisa8*) and L5a RSG (marked by *Hsd11b1* and *Gpr6*) cell types. In the future, researchers wishing to target cell types in other rat brain regions may benefit from verifying conservation of expression patterns, as done here for RSG.

### Transcriptomic data may reflect different excitation-inhibition balance

Our work identified a substantially larger total proportion of inhibitory neurons in rats as compared to mice, driven largely by an increase in the relative size of the Pvalb cluster. *In vivo*, *Pvalb*-positive interneurons are known to largely be fast-spiking (Kosaka et al., 1987) and play an important role in synchronizing high frequency oscillations such as gamma (Ahmed & Cash, 2013; Cardin et al., 2009; Hijazi et al., 2023; Hu et al., 2014; Sohal et al., 2009) Additionally, the balance between excitation and inhibition plays a critical role in neuronal information coding (Ahmed & Mehta, 2009; Denève & Machens, 2016; Ferguson & Gao, 2018; S. B. Nelson & Valakh, 2015; Tatti et al., 2017; Trevelyan & Watkinson, 2005; Zhou & Yu, 2018). Increased inhibitory control has been shown to be a feature of species with higher order cognitive functions such as humans and macaques, which have nearly three times the proportion of inhibitory neurons compared to mice (Loomba et al., 2022). If the increase in the proportion of inhibitory neurons in rats relative to mice is indicative of brain-wide increased inhibition, we then wonder whether it may play a role in any cognitive advantages which rats display over mice (Jaramillo & Zador, 2014).

In addition to establishing a strong foundation for the cross-species comparability of existing RSG literature, the present work offers itself as a guide to future electrophysiological, functional, and transcriptomic investigations of the RSG and RSC as a whole.

## ACKNOWLEDGEMENTS

This work was supported by NIH R34NS127101; NIH P50NS123067; Alzheimer’s Association Grant AARG-NTF-21-846572; NIH T32-DC000011 (CRK, TGE); NIH T32-DA007268 (TGE); NIH T32-NS076401 (CRK).

---

**Supplementary Figure 1.**
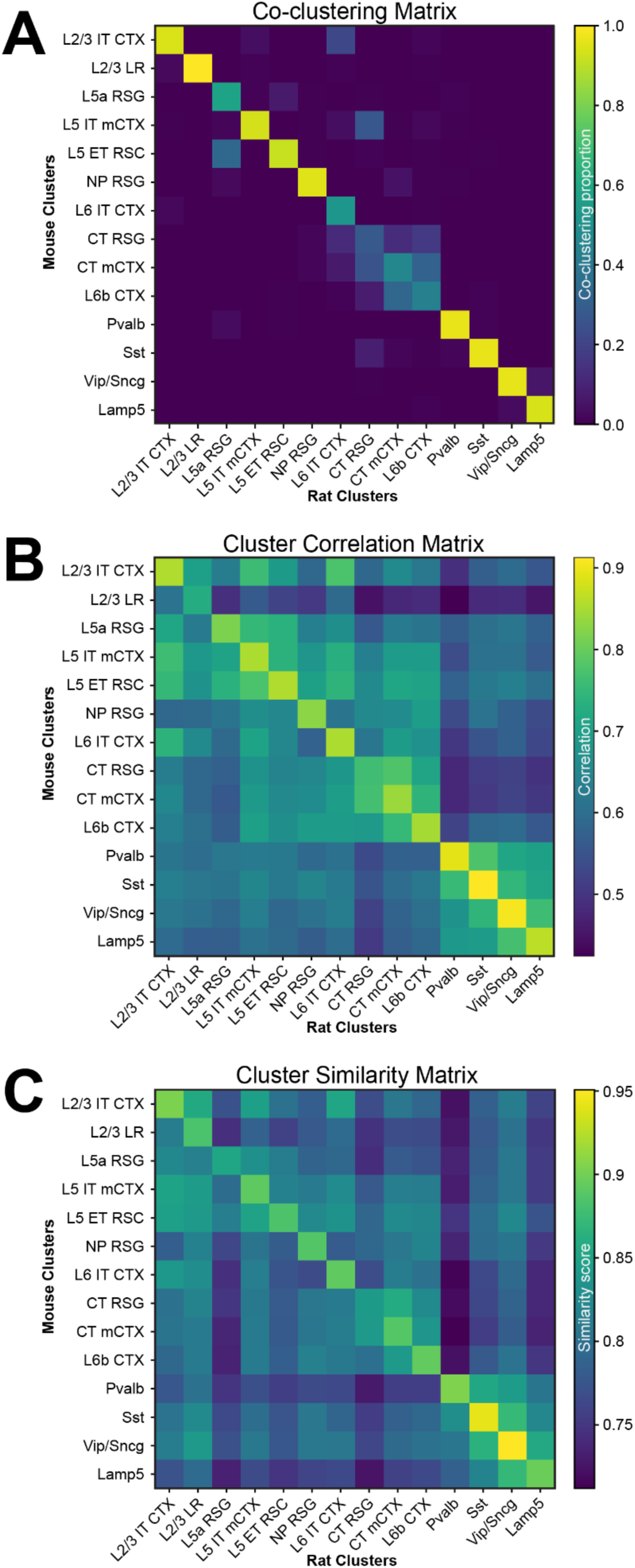
Additional methods of cross-species cluster alignment. See methods for details of how each matrix was computed. (A) Cross-species co-clustering matrix, normalized by the sum of each column, generated from 1000 random subsets of the mouse and rat datasets. Within each rat cluster (column), the value in the matrix represents the proportion of total co-clustering occurrences which came from a given mouse cluster. (B) Cross-species cluster correlation matrix computed from the mean log expression of highly variable genes. (C) Cluster similarity matrix computed from the ratio of mean intercluster and intracluster distances on each cross-species pair of clusters.

**Supplementary Figure 2.**
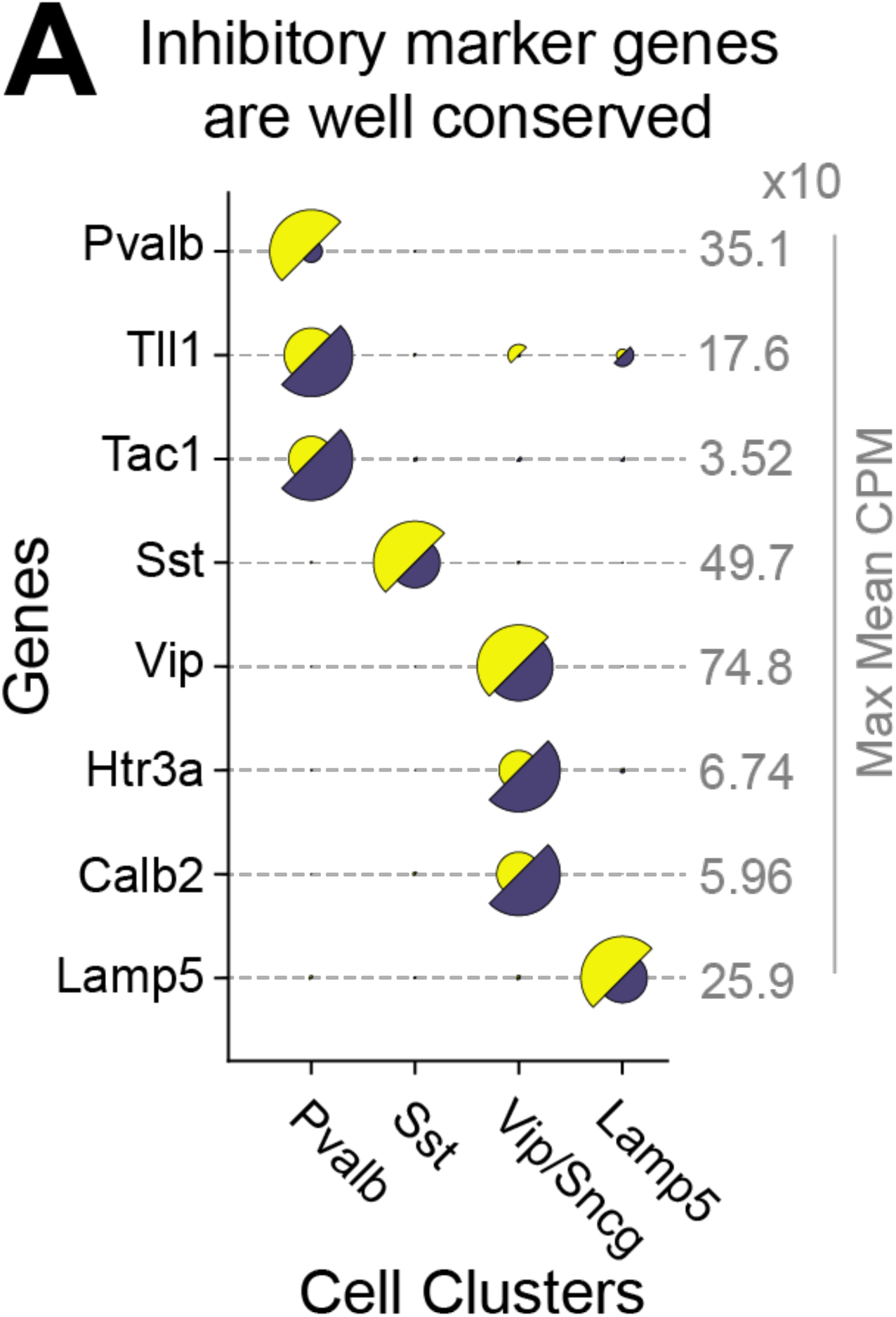
Inhibitory marker gene comparison between mouse and rat RSG. Half-circle plot of inhibitory marker genes demonstrating the selective and extremely well conserved nature of namesake genes *Pvalb*, *Sst*, *Vip*, and *Lamp5*, as well as selected others, in GABAergic clusters. Note that while the *Pvalb* gene exhibits a far lower **magnitude** of expression in rats, in this case this does not equate to a substantial change in the **proportion** of cells expressing: 81% of rat neurons in the Pvalb cluster express *Pvalb*, versus 91% of mouse neurons in the same cluster (percent expression data not shown).

**Supplementary Figure 3.**
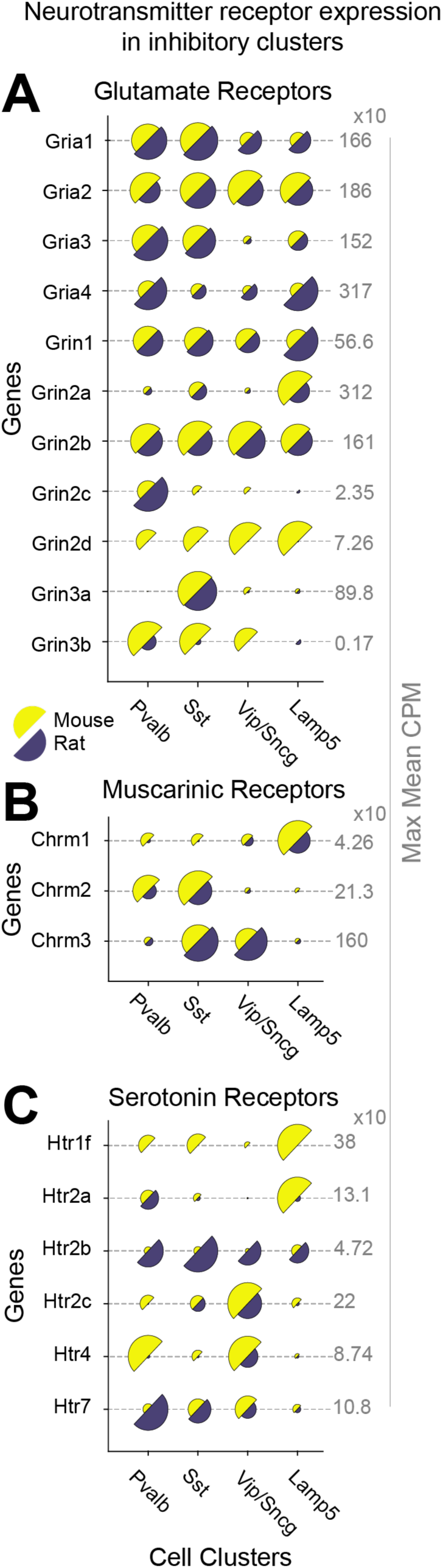
Neurotransmitter receptor expression in inhibitory clusters. (A to C) Half-circle plots of selected neurotransmitter receptor genes.

**Supplementary Figure 4.**
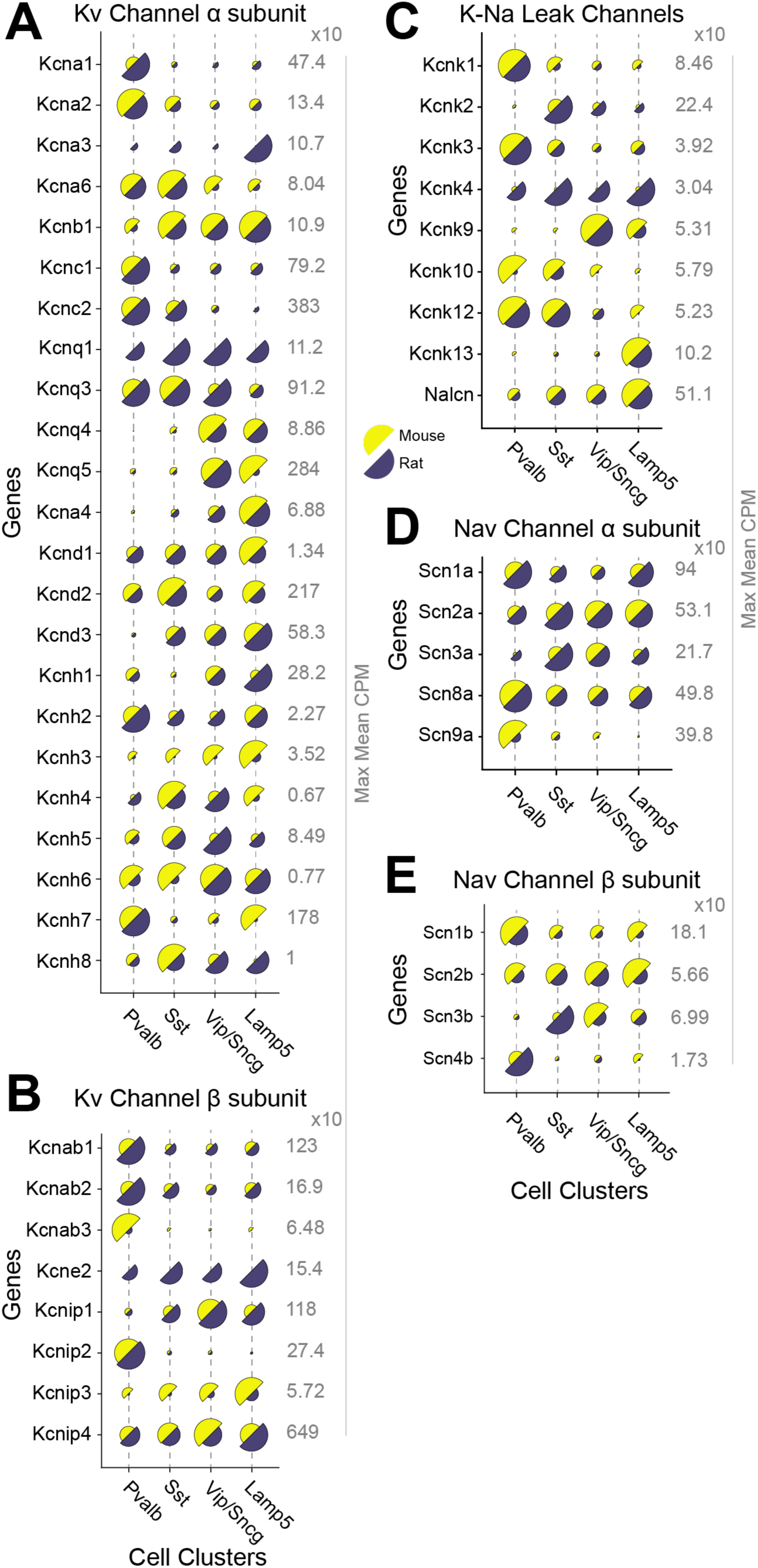
Ion channel expression in inhibitory clusters. (A to E) Half circle plots of selected ion channel gene expression.

## Notes

### Competing Interest Statement

The authors have declared no competing interest.

